# *CLN3* transcript complexity revealed by long-read RNA sequencing analysis

**DOI:** 10.1101/2023.10.12.562062

**Authors:** Hao-Yu Zhang, Christopher Minnis, Emil Gustavsson, Mina Ryten, Sara E Mole

## Abstract

**Background:** Batten disease is a group of rare inherited neurodegenerative diseases. Juvenile CLN3 disease is the most prevalent type, and the most common mutation shared by most patients is the “1-kb” deletion which removes two internal coding exons (7 and 8) in *CLN3*. Previously, we identified two transcripts in patient fibroblasts homozygous for the “1-kb” deletion: the “major” and “minor” transcripts. To understand the full variety of disease transcripts and their role in disease pathogenesis, it is necessary to first investigate *CLN3* transcription in “healthy” samples without juvenile CLN3 disease.

**Methods:** We leveraged PacBio long-read RNA sequencing datasets from ENCODE to investigate the full range of *CLN3* transcripts across various tissues and cell types in human control samples. Then we sought to validate their existence using data from different sources.

**Results:** We found that a readthrough gene affects the quantification and annotation of *CLN3.* After taking this into account, we detected over 100 novel *CLN3* transcripts, with no dominantly expressed *CLN3* transcript. The most abundant transcript has median usage of 42.9%. Surprisingly, the known disease-associated “major” transcripts are detected. Together, they have median usage of 1.51% across 22 samples. Furthermore, we identified 48 *CLN3* ORFs, of which 26 are novel. The predominant ORF that encodes the canonical CLN3 protein isoform has median usage of 66.7%, meaning around one-third of *CLN3* transcripts encode protein isoforms with different stretches of amino acids. The same ORFs could be found with alternative UTRs. Moreover, we were able to validate the translational potential of certain transcripts using public mass spectrometry data.

**Conclusion:** Overall, these findings provide valuable insights into the complexity of *CLN3* transcription, highlighting the importance of studying both canonical and non-canonical *CLN3* protein isoforms as well as the regulatory role of UTRs to fully comprehend the regulation and function(s) of *CLN3*. This knowledge is essential for investigating the impact of the "1-kb" deletion and rare mutations on *CLN3* transcription and disease pathogenesis.

## Background

The neuronal ceroid lipofuscinoses (NCLs, also known as Batten disease) are a group of rare inherited neurodegenerative lysosomal storage diseases characterised by the accumulation of autofluorescent lipofuscin and/or ceroid in lysosomes, with many causative genes. Juvenile CLN3 disease (Juvenile NCL, JNCL), is the most common, accounting for ∼50% of all NCLs cases (1). The first symptom of juvenile CLN3 disease is usually visual loss (retinitis pigmentosa), followed by seizures, cognitive and behavioural decline, motor impairment, and premature death in early adulthood (2). Classic juvenile CLN3 disease is an autosomal recessive disorder caused by mutations in *CLN3* gene (3). This gene encodes a 438 amino acids protein, most of which (amino acids 23-435) share similarities with members of the major facilitator superfamily (MFS) according to InterPro (4) (https://www.ebi.ac.uk/interpro/), though its function is not defined. Over 70% of patients with juvenile CLN3 disease are homozygous for a “1-kb” intragenic deletion (5), which is a deletion of 966- bp between intron 6 to intron 8 (rs1555468634, g.28485965_28486930del), causing the loss of two exons (coding exons 7 and 8) of the *CLN3* canonical transcript (5, 6). Previous research has identified at least two transcripts arising from the 1-kb deletion: a ‘major’ transcript, in which coding exon 6 is spliced to coding exon 9 out of frame so generating a premature stop codon in coding exon 9 with 28 novel amino acids; and a ‘minor’ transcript, in which coding exon 6 is spliced to coding exon 10 that brings the transcript back into the amino acid reading frame (7). These transcripts may exert different functions. Modelling the 1-kb deletion “minor” transcript in fission yeast suggests that it has some residual functionality as well as gaining novel function (8).

To date, there are six *CLN3* transcripts in Refseq (9) and 62 *CLN3* transcripts in Ensembl 110/GENCODE 44 (10, 11). Existing genome-wide Pacbio long-read RNA sequencing data in human and mouse cerebral cortex also suggests diversity in *CLN3* transcripts, with four novel *CLN3* transcripts identified in human data (12). The high variation in the numbers of different *CLN3* transcripts in current annotations raises the possibility of additional *CLN3* transcripts that have remained unannotated, as well as questions about which transcripts are functional.

To fully decipher the mechanism of juvenile CLN3 disease pathogenesis caused by the “1-kb” deletion, it is important to first understand *CLN3* transcription in “healthy” tissues. Given the existing complexity of *CLN3* transcription, several questions arise that need to be addressed: (a) How many *CLN3* transcripts remain unannotated? (b) How do *CLN3* transcripts vary, in terms of their open reading frames (ORFs) and untranslated regions (UTRs); (c) What is the expression of *CLN3* at both the transcript and gene level across tissues? As long-read RNA sequencing enables both the capture of full-length transcripts and their accurate quantification, these questions can now be addressed (13, 14). In this study, we addressed these questions by utilising the untargeted Pacbio long-read RNA sequencing datasets across 23 tissues and 10 cell lines available on the ENCODE portal (https://www.encodeproject.org) to study *CLN3* transcription.

## Methods

### Sequence similarity examination

Comprehensive gene annotation based on GENCODE release 29 (GENCODE 29) was downloaded from https://www.gencodegenes.org/human/release_29.html. All unique exons were studied, including 164 *CLN3* exons, and an additional 702,165 other exons. Exon sequences were extracted using the function getseq() of the R package BSgenome. Then BLASTN version 2.9.0 (15) was used with a threshold for filtering exons with a minimum percentage identity of 95% and a minimum bitscore of 100, compared to any given *CLN3* exons.

### GTEx V8 short-read RNA sequencing data

Gene-level quantifications in transcripts per million (TPMs), exon-exon junction read counts, and sample attributes of GTEx V8 were downloaded from GTEx portal (16) (https://gtexportal.org/home/datasets/) (GTEx_Analysis_2017-06-05_v8_RNASeQCv1.1.9_gene_reads.gct.gz, GTEx_Analysis_2017-06- 05_v8_STARv2.5.3a_junctions.gct.gz, GTEx_Analysis_v8_Annotations_SampleAttributesDS.txt).

Five exon-exon junctions which belong to the readthrough transcripts, i.e., splice donors in *CLN3* and splice acceptors in *NPIPB7,* were selected and investigated: 1) chr16:28466903-28476250:- as shown in transcript ENST00000635887; 2) chr16:28466903-28482104:- as shown in transcript ENST00000637378; 3) chr16:28466903-28477463:- as shown in transcripts ENST00000637376, ENST00000636078, ENST00000637745, ENST00000568224 and ENST00000636503; 4) chr16:28471175-28476250:- as shown in transcripts ENST00000636017, ENST00000636866, ENST00000637299, and ENST00000638036; and 5) chr16:28466903-28477011:- as shown in transcript ENST00000636766.

The detection rates of these junctions were determined by proportion of tissue donors in which a specific exon-exon junction was detected. For tissues with 100 or more donors, tissues with the maximum detection rates for specific junctions were checked. For each tissue, the average read counts for specific junctions across all donors were calculated. Then the average read count for each junction was investigated for minimum, mean, and maximum values across all tissue types.

### Long-read RNA sequencing data processing

#### ENCODE untargeted long-read RNA sequencing data

ENCODE untargeted long-read RNA sequencing data was downloaded from the ENCODE portal (17) (https://www.encodeproject.org/). We selected 99 samples from nine ENCODE-defined organs, including blood vessel, brain, connective tissue, embryo, endocrine gland, heart, blood, lung, and skin. Data from cancer cell lines was not included. The processed quantification files in .tsv format and annotation files in .gtf format were downloaded for further analysis. All the downloaded quantification and annotation files were generated by ENCODE long-read RNA sequencing pipeline (<u>GitHub - ENCODE-DCC/long-read-rna-pipeline: ENCODE long read RNA-seq pipeline</u>). Corrected transcripts were annotated and quantified by the TALON package (18) (https://github.com/mortazavilab/TALON). GENCODE 29 was used as a reference when generating these datasets.

#### Transcript separation

For all downloaded ENCODE annotation files, transcripts overlapping with the *CLN3* locus (chr16:28474111-28495575:-) were selected using GffRead (19). To separate *CLN3* transcripts and transcripts of the readthrough gene ENSG00000261832, transcripts that also overlapped with the *NPIPB7* locus (chr16:28456329-28472336:-) were identified. Transcripts that overlapped with both *CLN3* and *NPIPB7* loci were assigned to ENSG00000261832. Transcripts that overlapped with the *CLN3* locus but not with the *NPIPB7* locus were assigned to *CLN3*.

#### Gene-level quantification

The quantification of transcripts in read counts within each sample was obtained from downloaded quantification files. All transcripts marked as “Genomic”, and transcripts of genes marked “Antisense” or “Intergenic” were removed. The read counts for each gene were calculated by summing the read counts of all transcripts identified at each locus. Then for each sample, TPM for each gene was calculated by: 1) calculating the reads per kilobase (RPK) for each gene by dividing the read count by the length of gene in kilobases; 2) calculating the scaling factor by dividing the sum of RPK values within the sample by 1,000,000; 3) calculating the TPM for each gene by dividing the RPK value for each gene by the scaling factor. Then the TPMs of *CLN3* and ENSG00000261832 were checked within each ENCODE-defined organ.

#### Open reading frame (ORF) prediction

The ORFs for *CLN3* transcripts were predicted using the findMapORFs() function from the R package ORFik (20) (https://github.com/Roleren/ORFik). Exon coordinates were obtained from annotation files downloaded from ENCODE portal. GRCh38 was used as the reference genome. “ATG” and “TAA|TAG|TGA” were used as the start codon and stop codons, respectively. When predicting the ORFs, only the longest ORF per stop codon was kept. Then, the longest ORF per transcript was selected and the length of the product was calculated. ORF identifiers (ID) were assigned to each ORF, for example, “CLN3_24_438aa”, based on the length rank (e.g., ‘24’) and length of the product (e.g., 438 amino acids).

#### Transcript merging

Transcripts were called separately in each sample by the ENCODE pipeline so to study all transcripts and their specific features using a common framework and naming convention, transcripts were merged based on the following criteria: (a) the same ORF; (b) the same proximal 5’ and 3’UTR internal boundaries; (c) distal 5’ and 3’UTR ends located within 20 bps.

UTRs were extracted using the fiveUTRsByTranscript() and threeUTRsByTranscript() functions from R package GenomicFeatures (21). The same ID was assigned for a given UTR with the same internal boundaries and with 5’ and 3’ ends within 20 bps (+/-). IDs for UTRs (e.g., “5UTR_136”) consisted of type (e.g., ‘5’ or ‘3’) and arbitrary numbers (e.g., ‘136’).

For the merged transcripts, IDs were assigned using a combination of ORF IDs, 5’UTR IDs, and 3’UTR IDs. For example, "CLN3_125_316aa_5UTR_132_3UTR_79". This allows an immediate appreciation of the structure which would not be possible using a completely arbitrarily assigned ID. Transcripts detected in three or more samples were considered valid.

#### Transcript-level quantification

For merged transcripts, transcript-level expression was assessed on the basis of transcript occurrence and usage. Transcript occurrence refers to the number of samples in which a specific transcript was detected. Transcript usage refers to all transcription that was assigned to a specific *CLN3* transcript divided by the total transcription from the locus (in read counts). Tissue-specific transcripts were defined as transcripts that had usage in a given tissue that was at least two times higher than in any other tissue.

#### Nonsense-mediated decay prediction

We predicted whether a transcript was subject to nonsense-mediated decay (NMD) using the function predictNMD() from R package factR (22) (https://github.com/fursham-h/factR). This function is based on the commonly used rule that if the stop codon of a transcript is more than 50 nucleotides upstream of the most downstream exon-exon junction, the transcript will be NMD- sensitive.

#### Determining the novelty of ORFs, UTRs and transcripts

Known *CLN3* transcripts were extracted from GENCODE version 29. Then, ORFs were predicted by R package ORFik (20) using the same arguments as above. UTRs from GENCODE 29 and ENCODE datasets were extracted using fiveUTRsByTranscript() and threeUTRsByTranscript() functions from R package GenomicFeatures. ORFs/UTRs detected in GENCODE 29 but not in the selected ENCODE datasets were considered novel. Merged transcripts were classified as novel if all the ENCODE transcripts they were derived from were novel.

#### Transcripts classification

Transcripts were categorised into different types based on NMD prediction, coding potential and novelty. Transcripts that were predicted to undergo NMD were categorised based on their novelty into NMD_Known and NMD_Novel. Non-NMD transcripts with ORFs encoding products smaller than 150aa were considered non-coding. These non-coding transcripts were categorised into Non_coding_Known and Non_coding_Novel based on their novelty; and then coding transcripts were categorised into Coding_Known and novel coding transcripts. Novel coding transcripts were further categorised based on the novelty of the UTRs and ORFs: transcripts with novel 3’ or 5’ UTRs but known ORFs (Novel_3’/5’UTR_only), coding transcripts with both known ORFs and known UTRs but the combination of UTRs and ORFs were novel (Novel_combination), coding transcripts with novel ORFs and at least one novel UTR (Novel_ORF_and_UTR), and coding transcripts with only novel ORFs (Novel_ORF_only).

#### Transcripts visualisation

*CLN3* transcripts were visualised using R Package ggtranscript (23) (https://github.com/dzhang32/ggtranscript). The coding sequences (CDSs) coordinates used were generated by the ORF prediction step using the R package ORFik.

#### Validation of transcripts

Transcripts were validated in terms of their 5’ transcription start sites (TSSs), splice junctions, and 3’ polyadenylation sites (PASs). The latest reference TSS (refTSS) data in BED format was downloaded from http://reftss.clst.riken.jp/. This included 5’ sequencing data from FANTOM5, EPDnew, ENCODE RAMPAGE, ENCODE CAGE, stem cell CAGE (DDBJ accession number DRA000914), and dbTSS. The integrated data was reprocessed and mapped to the latest version of the reference genome (24). RJunBase (www.RJunBase.org) is a web database of splicing junctions from 18084 normal samples and 11540 cancer samples (25). A total number of 64 non-tumour-specific linear splice junctions of *CLN3* were identified and downloaded. Poly-Adenylation annotation for human GRCh38.96, Homo sapiens v2.0, was downloaded from PolyAsite 2.0 (https://polyasite.unibas.ch/atlas#2). The polyA human atlas included 221 different 3’end sequencing libraries prepared by different protocols, including 3’-Seq (Mayr), 3’READS, DRS, QuantSeq_REV, SAPAS, PAPERCLIP, PolyA-seq, PAS-seq, A-seq, and 3P-Seq (26).

### Public mass spectrometry data

Public mass spectrometry datasets PXD026370 and PXD028605 were downloaded from ProteomeXchange (https://www.proteomexchange.org/). PXD026470 contains data derived from post-mortem human brain tissue from patients with multiple system atrophy (MSA) (N = 45) and controls (N = 30) (27). PXD028605 contains data from non-small-cell lung cancer (NSCLC) patients (N = 5) and healthy individuals (N = 5) whole blood cell pellets, and whole blood collected from healthy volunteers (28).

Transcripts predicted to have ORFs with unique peptide sequences were selected. In each case, the public mass spectrometry datasets were searched using MetaMorpheus (29) 1.0.1 for evidence of the unique peptide sequence with default settings to determine their presence. Alignments of identified peptides and corresponding CLN3 protein sequences were visualised using R package ggmsa (30).

### Protein structures

Translated amino acid sequences of transcripts of interest were used as input for AlphaFold 2.3.2 Colab to enable prediction of the protein structure (31) (https://colab.research.google.com/github/deepmind/alphafold/blob/main/notebooks/AlphaFold.ipynb). Pairwise Structure Alignment was performed using jFATCAT (rigid) on RCSB Protein Data Bank (32) (https://www.rcsb.org/).

## Results

### *CLN3* transcription shows complexity in current annotation

Of the 62 *CLN3* transcripts in Ensembl 110, 29 of them are classified as protein-coding, and 10 of them have the highest transcript support level with all splice junctions supported by mRNAs from other sources (e.g., SPTrEMBL and RefSeq). Currently, only 172 of the 23,217 protein-coding genes in Ensembl 110 have more than 50 transcripts, and *CLN3* is ranked 81^st^ among all protein-coding genes in terms of number of transcripts (**Fig S1)**.

### *CLN3* overlaps with readthrough gene ENSG00000261832

Multimapping of short-read RNA-sequencing data can cause inaccuracies in transcript annotation and quantification when reads cannot be uniquely assigned and are instead discarded (33). To assess whether multimapping could be limiting the annotation of *CLN3* transcripts we first determined if there was any other gene with high sequence similarity to the locus. The sequence information for all 164 annotated exons of *CLN3* was aligned against all other exons in GENCODE 29. This analysis demonstrated that 42 *CLN3* exons are identical in sequence to exons of ENSG00000261832, a readthrough gene containing exons from both *CLN3* and *NPIPB7.* All other *CLN3* exons were identically assigned to the *CLN3* locus alone. To examine the expression of this *CLN3*-*NPIPB7* readthrough gene across human tissues, we used data generated by GTEx V8 (16) (https://gtexportal.org/home/datasets/). Since ENSG00000261832 was not included in gene-level quantifications, we analysed the expression of five unique exon-exon junctions of ENSG00000261832 transcripts (splice donor sites within *CLN3* and splice acceptor sites within *NPIPB7,* **Fig 1**) using GTEx V8 exon-exon junction read counts data. All five exon-exon junctions are detected in multiple tissues, supporting their existence. Amongst all five junctions, “junction 3” (chr16:28466903-28477463:-) has the highest detection rate at 95.3% (364 tissue donors out of 372) in brain samples and the highest average read counts of 2.12 across all tissues (**Table S1**). “Junction 2” (chr16:28466903-28482104), which is specific to the transcript (ENST00000637378) with an open reading frame containing coding sequences from both *CLN3* and *NPIPB7,* is detected in 63.2% of brain tissue donors (**Table S1**).

**Figure 1.**
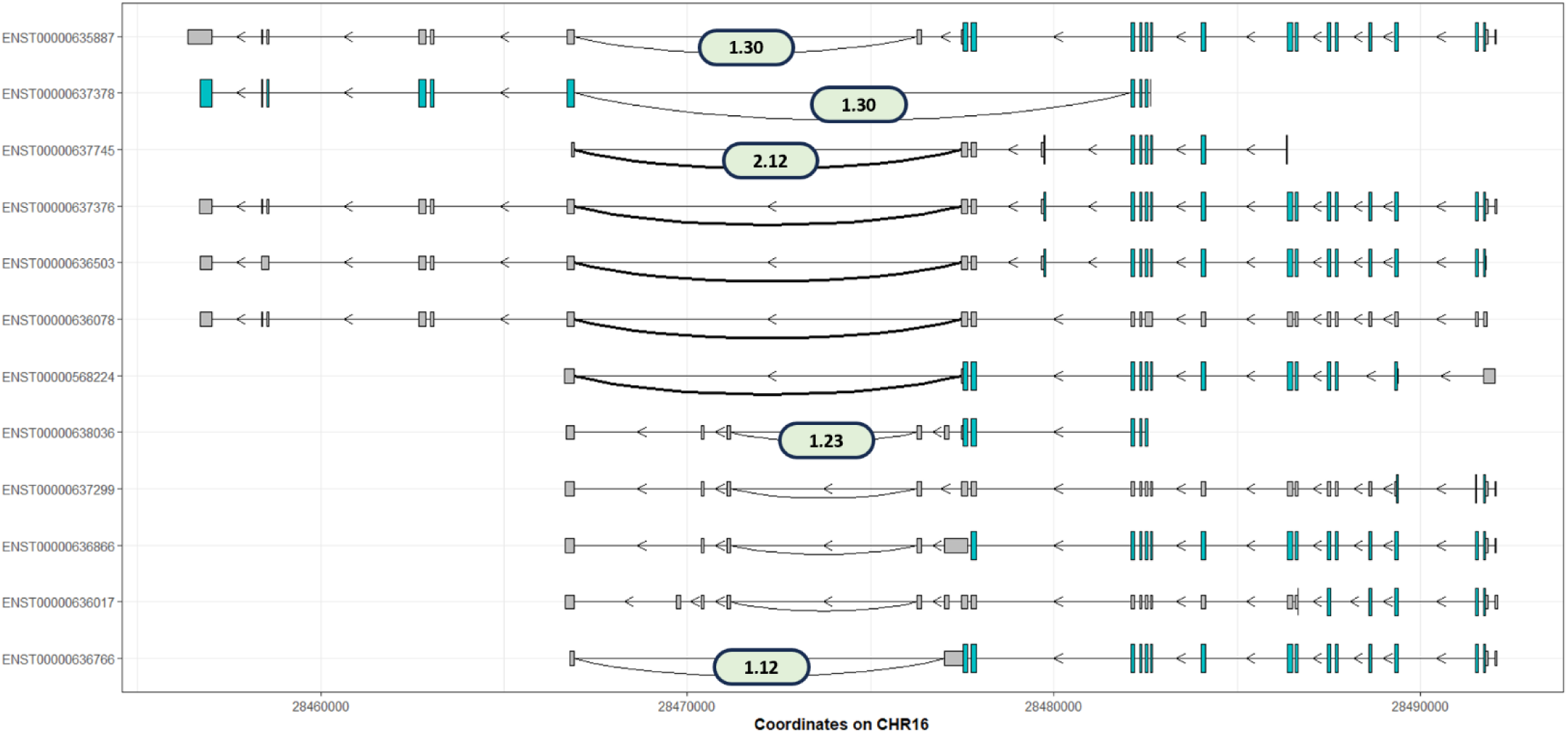
Summary of exon-exon junctions specific to the readthrough gene ENSG00000261832 from GTEx V8. All five exon-exon junctions with splice donor sites in *CLN3* locus and splice acceptor sites in *NPIPB7* locus are extracted from GTEx V8 exon-exon junction read counts data. All 12 transcripts containing these junctions are plotted, with taller coloured boxes showing the open reading frames (ORFs) and shorter grey boxes showing the untranslated regions (UTRs). Note, these transcripts are present on the antisense strand, so read right to left. These five junctions are shown in curved lines connecting the donor and acceptor sites. For each junction, the average read count among all tissue donors and tissue types is shown in boxes above the junction.

### *CLN3* annotation is complicated by the *CLN3-NPIPB7* readthrough gene

To examine the accuracy of *CLN3* annotation and quantification, we analysed ENCODE long-read RNA-sequencing data. Transcripts that overlap with the *CLN3* locus (chr16:28474111-28495587:-) were selected within 99 chosen samples, which includes data generated from 23 different human tissue types and 10 different human cell lines (**Table S2**). The selected transcripts were annotated to either *CLN3* or the *CLN3*-*NPIPB7* readthrough gene, ENSG00000261832 (**Fig 2A and B**). To avoid mis-annotation, we categorised selected transcripts that overlap with the *CLN3* locus based on whether they also overlap with the *NPIPB7 locus*. Those transcripts that overlap with both *CLN3* and *NPIPB7* loci were assigned to ENSG00000261832. We found that ∼10-25% of all *CLN3* transcripts had originally been assigned to the readthrough gene, ENSG00000261832 by the ENCODE Long Read RNA-Seq Analysis Protocol for Human Samples (**Fig 2C**).

**Figure 2.**
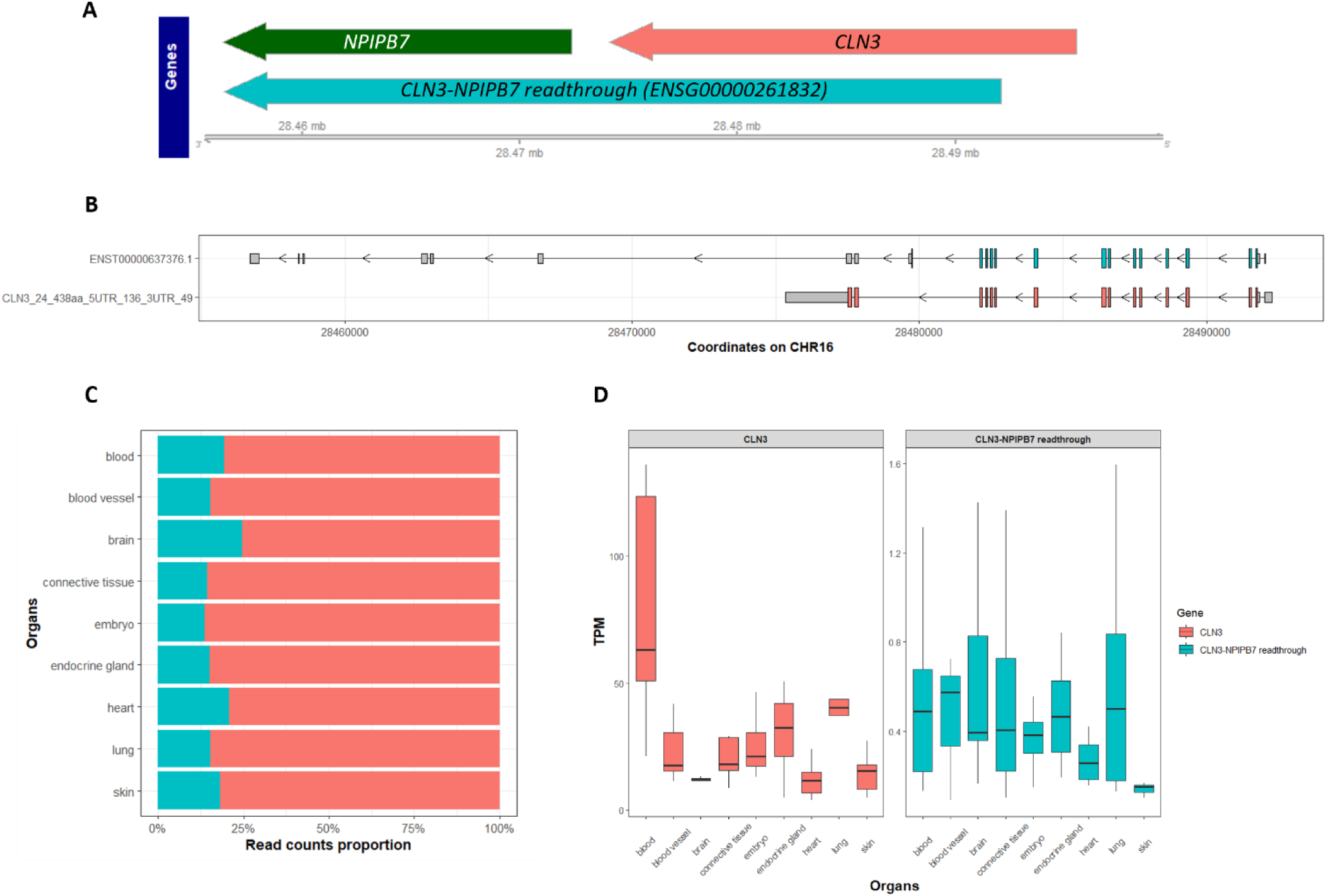
Assigning *CLN3* transcripts to the *CLN3-NPIPB7* readthrough gene affects the accurate quantification of *CLN3* in ENCODE long-read RNA sequencing data. **A** The *CLN3, NPIPB7,* and *CLN3*-*NPIPB7* readthrough gene (ENSG00000261832) loci are shown according to their genomic coordinates from Ensembl 110. **B** The readthrough gene, ENSG00000261832, contains exons of *CLN3* and *NPIPB7.* This plot shows representative transcripts of *CLN3* and ENSG00000261832. Coding sequences are coloured by red for *CLN3* and cyan for ENSG00000261832, grey boxes show the UTRs. Note, *CLN3* transcript is present on the antisense strand, so it reads right to left. **C** Over 10% of *CLN3* transcripts were assigned to the readthrough gene in selected ENCODE data, affecting the accurate annotation and gene-level quantification of *CLN3.* D The gene-level quantification of *CLN3* and the readthrough gene ENSG00000261832 across nine ENCODE-defined organs are plotted. The readthrough gene has rather lower TPMs across all samples compared with *CLN3*.

*CLN3* is expressed in all 99 samples. We observed that gene-level expression of *CLN3* is variable across different organs (**Table 1**). Among these selected organs, *CLN3* has the highest expression in blood (median TPM 63.10), and the lowest expression in heart (median TPM 11.59). Compared with GTEx V8 bulk gene expression for *CLN3,* our data shows similar variability across different organs. Both datasets show relatively low expressions in the brain and heart, and higher expressions in blood and lung (**Table 1**). In contrast, the readthrough gene, ENSG00000261832, is not detected in all samples of a given tissue, for example, it is only detected in 35.3% (6 out of 17) of heart samples. It exhibits low expression levels across all nine organs, with median TPMs ranging from 0.15 (skin) to 0.57 (blood vessel) (**Fig 2D, Table S3**).

**Table 1.** Gene-level quantification of *CLN3* across nine ENCODE-defined organs.

| Organs | Number of Samples | Tissues | Min TPM | Median TPM | Max TPM | GTEx V8 median TPMs within specific tissues |
| --- | --- | --- | --- | --- | --- | --- |
| Blood | 16 | HL-60 | 21.20 | 63.10 | 136.10 | 19.07 |
| Blood vessel | 6 | posterior vena cava, aorta, endothelial cell of umbilical vein | 11.44 | 17.56 | 41.87 | 11.52 – 13.36 |
| Brain | 9 | middle frontal area 46 | 5.54 | 11.97 | 13.25 | 3.31 - 10.47 |
| Connective tissue | 16 | IMR-90, osteocyte, WTC11, chondrocyte, mesenteric fat pad, HFFc6 | 8.61 | 18.07 | 62.77 | NA |
| Embryo | 18 | H9, neural crest cell, endodermal cell, H1 | 12.93 | 21.09 | 46.38 | NA |
| Endocrine gland | 9 | adrenal gland, type B pancreatic cell, right lobe of liver, progenitor cell of endocrine pancreas | 4.93 | 32.51 | 50.77 | 6.10 – 16.76 |
| Heart | 17 | heart left ventricle, heart right ventricle, left cardiac atrium, right cardiac atrium, cardiac septum, Right ventricle myocardium inferior, left ventricle myocardium superior, left ventricle myocardium inferior, Right ventricle myocardium superior | 4.02 | 11.59 | 24.08 | 3.20 – 4.55 |
| Lung | 6 | upper lobe of right lung, lower lobe of left lung, lower lobe of right lung, left lung | 23.25 | 40.25 | 107.08 | 22.18 |
| Skin | 8 | WTC11, GM23338 | 4.79 | 15.50 | 27.08 | 16.61 - 17.85 |

### 172 different *CLN3* transcripts are detected with no dominantly expressed transcript

We identified 172 *CLN3* transcripts across 99 samples. They were categorised based on their novelty, likelihood of undergoing nonsense-mediated decay (NMD) and coding potential (**Fig 3A**). Among these 172 transcripts, 147 are absent from GENCODE 29, including 75 transcripts with coding potential. Within these 75 novel coding transcripts, 13 have novel ORFs and known UTRs; 34 have novel ORFs and novel 5’ or 3’UTRs; 25 have only novel 5’UTRs; and 3 have known ORFs and UTRs but novel combinations (**Fig 3B**).

**Figure 3.**
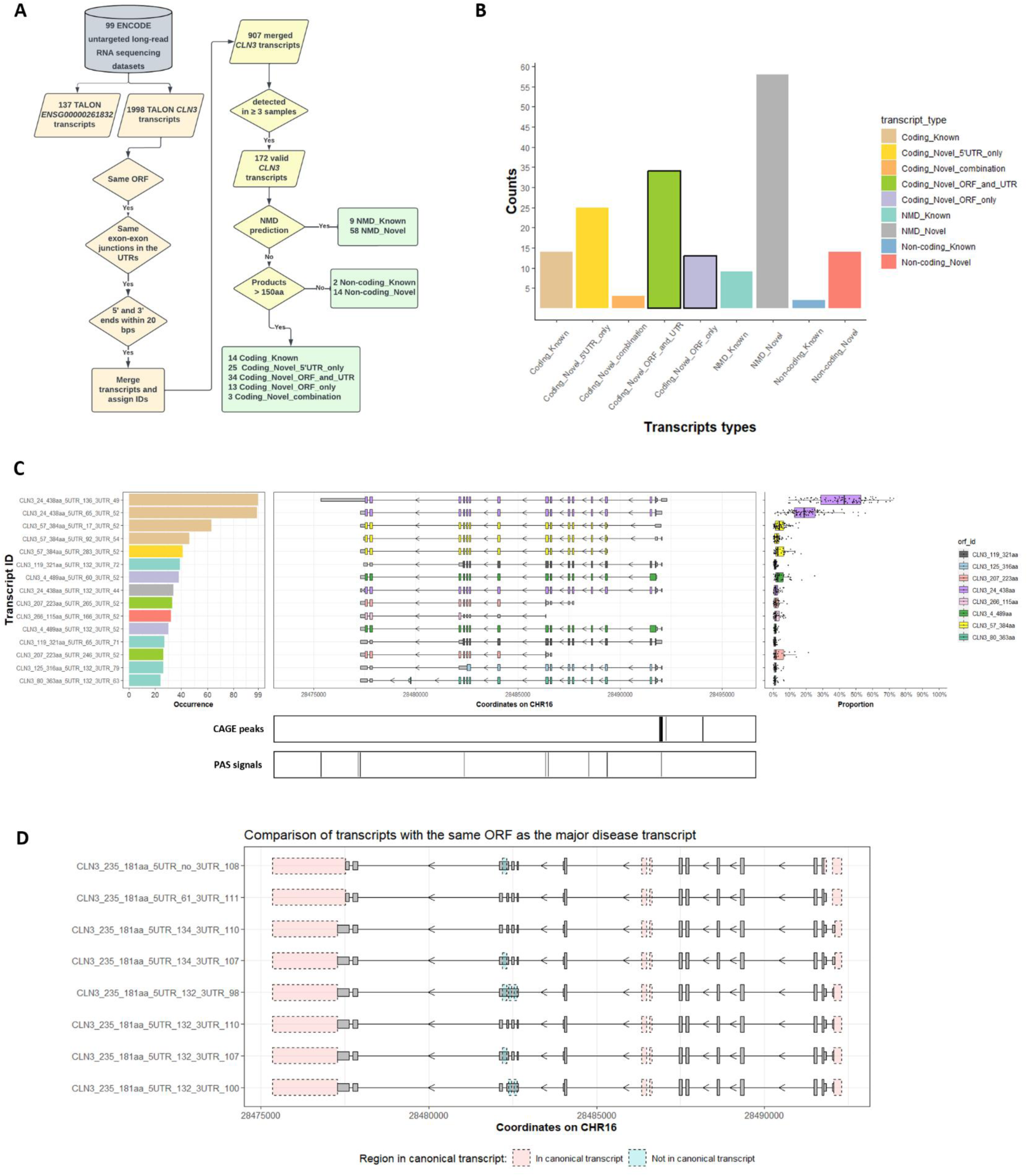
Summary of identified *CLN3* transcripts. **A** Data processing pipeline for ENCODE long-read RNA sequencing data is shown. **B** Valid transcripts are categorised based on transcript novelty, ORF and UTR novelty, coding potential and NMD prediction, numbers of transcripts within different categories are shown in the bar plot. Bars showing the number of transcripts with novel ORFs are highlighted. **C** Top 15 *CLN3* transcripts based on the number of samples in which they are detected (occurrence, left panel) are selected. The occurrence numbers are shown in the left panel, coloured by categories of these transcripts. The middle panel shows the structures of transcripts, the coloured taller boxes show the ORFs while the grey shorter boxes show the UTRs. The transcript usage is shown in the right panel. Note, CLN3 transcripts are present on the antisense strand, so read right to left. TSSs and PASs signals are shown under the transcript structures, aligned by genomic coordinates. **D** Eight transcripts with the same ORF as the “major” disease-associated transcript but different UTRs are detected. This plot shows the structural differences between these “major” transcripts and the canonical *CLN3* transcript “CLN3_24_438aa_5UTR_136_3UTR_49”. All grey boxes show structures of the “major” transcripts, with taller grey boxes showing the ORF and shorter grey boxes showing the UTRs. The pink boxes show sequences in the canonical transcript but not in the “major” transcripts, blue boxes show sequences in the “major” transcripts but not in the canonical transcript.

The expression of transcripts was analysed, both in terms of the number of samples in which a transcript is detected and the usage of that transcript across tissues. Interestingly, unlike previous research showing that dominant transcripts of protein-coding genes account for around 80% of transcription of the locus (34), there is no dominant transcript detected for *CLN3*. The most abundant transcript is the canonical *CLN3* transcript “CLN3_24_438aa_5UTR_136_3UTR_49” as in GENCODE 29, which is detected in all 99 samples and has median usage of 42.9%. The second most abundant transcript “CLN3_24_438aa_5UTR_65_3UTR_52” has the same ORF of 438aa but different UTRs and is detected in almost all samples (N = 98), with median usage of 18.7%. The third most abundant transcript, “CLN3_57_384aa_5UTR_17_3UTR_52”, has a smaller ORF due to a non-canonical start codon; this is detected in 63 samples with median usage of 3.7% (**Fig 3C)**.

Transcripts can show tissue-specific usage (35). In this study, tissue-specific transcripts are defined as those exhibiting 2-fold higher expression in one tissue as compared to that in any other tissue. Amongst the 172 *CLN3* transcripts, 75 tissue-specific transcripts are detected (**Table S3, Fig S2**). Across nine organs, blood contains the largest number of tissue-specific transcripts (N = 21), followed by heart (N = 15) and embryo (N = 10) (**Table S4**).

### “Disease-associated” transcripts are detected in control samples

We identified transcripts matching the splicing patterns of those previously reported in patient-derived fibroblasts homozygous for the 1-kb deletion namely the “major” and “minor” disease-associated transcripts (7). In this dataset, we identified eight transcripts showing the same ORF pattern as the “major” disease-associated transcript, that is coding exon 6 spliced to exon 9 introducing a non-canonical coding sequence and premature stop codon; these transcripts have the same ORF but different 5’/3’ UTRs (**Fig 3D**). Together, they have median usage of 1.51% and are detected in 22 samples from 13 tissue donors. We also identified two transcripts showing coding exon 6 spliced to coding exon 10 and the canonical stop codon as the “minor” disease-associated transcript, but these transcripts did not pass the filtration of detection in three or more samples. These two “minor” transcripts are detected once in male newborn endothelial of umbilical vein primary cell and female embryo (5 days) chondrocyte *in vitro* differentiated cells, respectively.

### 48 different ORFs are detected across all *CLN3* transcripts

Different transcripts can contain the same open reading frames (ORFs) and produce the same protein products. Therefore, to infer the proportion of different CLN3 protein isoforms generated, we investigated the usage of ORFs. ORF usage was calculated by summing all transcripts containing the same ORFs in each individual sample. From the 99 selected samples, we detected 48 different ORFs, of which 26 are novel. The most abundant ORF is that encoding the canonical CLN3 protein isoform of 438aa (UniProt ID: Q13286-1), “CLN3_24_438aa”. Summing all transcripts containing this ORF gave median usage of 66.7% (**Fig 4A**), suggesting that transcripts encoding non-canonical protein isoforms account for around one-third of *CLN3* transcription. The most abundant non-canonical ORF is “CLN3_57_384aa” which encodes a shorter protein of 384aa (UniProt ID: B4DFF3) generated through a different start codon position that misses the first two coding exons present in the canonical transcript. Collectively, transcripts containing the “CLN3_57_384aa” ORF have median usage of 7.4% (**Fig 4A**) and are detected in 87 samples. The most abundant novel ORF is “CLN3_207_223aa” which has a start codon in the coding exon 8 of the canonical transcript and the canonical stop codon. It has median usage of 3.1% (**Fig 4A**) and is detected in 72 samples.

**Figure 4.**
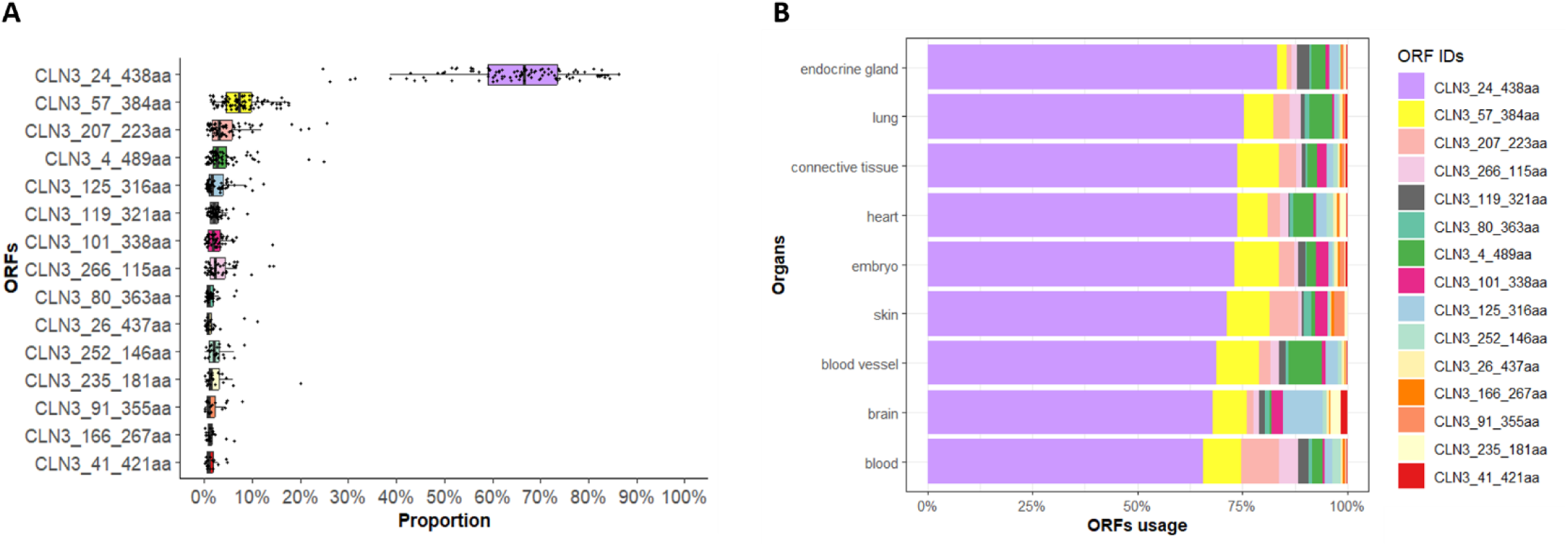
Summary of the top 15 CLN3 ORFs. The top 15 ORFs calculated by summing usage of all *CLN3* transcripts containing specific ORFs are plotted in **A**; the canonical *CLN3* ORF “CLN3_24_438aa” has a median usage of 66.7%, therefore around one-third of *CLN3* transcripts encode different protein isoforms. The top 15 ORFs’ usage in different organs is plotted in **B**. The usage of the transcripts containing non-canonical ORFs is variable across organs.

Additionally, the ORF identified in the “major” disease-associated transcript, “CLN3_235_181aa” (UniProt ID: Q9UBD8), has median usage of 1.51% across 22 samples. Moreover, the same ORFs could have different usage across different organs. For example, the ORF “CLN3_125_316aa” which is generated through a retained intron event and a different stop codon (**Fig 5A**) is highly expressed in brain while “CLN3_207_223aa” is highly expressed in blood. These particular transcripts encode protein isoforms with different stretches of amino acid sequences and so may have different functional properties.

**Figure 5.**
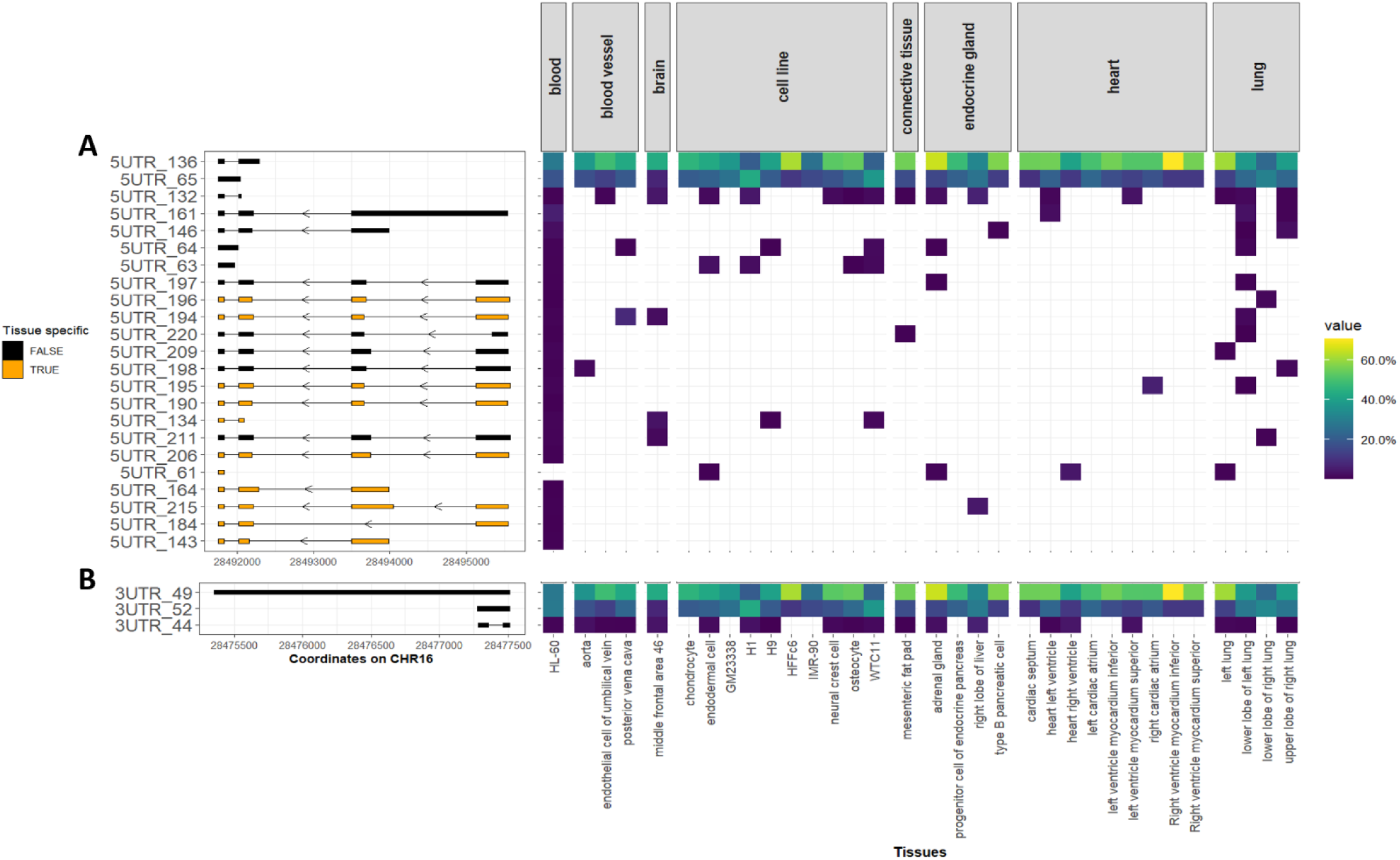
The same CLN3 ORFs are associated with different 5’UTRs and 3’UTRs. The presence of UTRs for three selected CLN3 ORFs is plotted; UTRs with tissue-specific expression are shown in orange, and non-tissue-specific UTRs are shown in black. The usage of these UTRs (with specific ORFs) in different tissues is plotted using heat maps with yellow showing high usage and dark blue showing low usage. Cell lines are grouped together and shown separately. **A** Twenty-three different 5’UTRs were found with the canonical *CLN3* ORF “CLN3_24_438aa”, 16 of them show additional upstream exons, 11 of them show tissue-specificity; **B** Three different 3’UTRs were identified with the canonical ORF “CLN3_24_438aa”, none of them shows tissue-specificity. The distal 3’UTR, “3UTR_49”, is specific to the canonical ORF.

### *CLN3 t*ranscripts with the same ORFs have alternative UTRs

Among the 172 *CLN3* transcripts, 82 different 5’UTRs were identified, 69 of which are novel. Twenty-nine of the 48 ORFs were found with multiple 5’UTRs. The canonical ORF “CLN3_24_438aa” has the most 5’UTRs (N = 23) (**Fig 5A**). Out of the 23 5’UTRs found with “CLN3_24_438aa,” 16 of them show distal transcription start sites (TSSs) and additional exons at the 5’ ends compared to the 5’UTR of the canonical transcript, i.e., “5UTR_161”. By examining the usage of transcripts with specific ORFs and specific 5’ or 3’ UTRs, we found that for ORF “CLN3_24_438aa”, there are 11 5’UTRs showing tissue-specific patterns, with blood harbouring the most tissue-specific 5’UTRs (N = 5). Out of these tissue-specific 5’UTRs, “5UTR_194” has the highest usage of 8.3% in blood vessel.

There are fewer 3’UTRs than 5’UTRs. Thirty-one different 3’UTRs were identified, of which 23 are novel. Only 12 ORFs were found with multiple 3’UTRs. There are three 3’UTRs found with ORF “CLN3_24_438aa” (**Fig 5B**). The distal 3’UTR, “3UTR_49”, is the only distal 3’UTR identified within 172 *CLN3* transcripts. It is specifically detected with the canonical ORF.

### The variety of *CLN3* transcripts can be validated

Given the large variety of *CLN3* transcripts identified, we sought to validate these transcripts and their translation potential using data from different independent technologies. The transcription start sites (TSSs) of *CLN3* transcripts, which represent 5’ ends of these transcripts, were validated using the reference TSS dataset from refTSS (24). Among the 172 *CLN3* transcripts, TSSs of 65.7% of them (N = 113) are located within 50bps of the TSS peaks, providing confidence in the existence of these transcripts. There are TSSs signals upstream of the canonical *CLN3* transcript which indicate that *CLN3* might be controlled by upstream promoters (**Figure 3C**).

We found that 41 out of 58 splice junctions found in *CLN3* transcripts are detected in RJunBase (www.RJunBase.org). The absence of 17 splice junctions suggests that the annotation of *CLN3* is not complete.

The polyadenylation (polyA) sites, which represent the 3’ ends of transcripts, of 91.3% of *CLN3* transcripts (N = 157) match with the polyA sites from polyAsite 2.0 (26). PolyA signals of the distal 3’UTR, “3UTR_49”, and shorter 3’UTRs, for example, “3UTR_52” are detected in polyAsite 2.0 data (**Figure 3C**). Strong polyA signals are also found in the region of coding exons 7 and 8, and exon 3 of the canonical CLN3 transcript (**Figure 3C**). These polyA signals could indicate premature termination of transcription. However, these premature transcription termination sites are not detected in our selected samples.

### The translation of *CLN3* transcripts can be validated

Finally, we investigated the translational potential of transcripts predicted to have an ORF (including NMD transcripts) using public mass spectrometry data from human post-mortem brain and whole blood. Unique sequence stretches from seven specific ORFs were tested: CLN3_4_489aa (retained intron), CLN3_80_363aa (retained intron), CLN3_125_316aa (retained intron), CLN3_48_414aa (exon skipping), CLN3_101_338aa (alternative translation start site, exon skipping), CLN3_238_181aa (“major” disease-associated transcript, exon skipping), CLN3_115_328aa (“minor” disease-associated transcript, exon skipping) (**Figure 6A**). Peptides “ADSAPGGHARSGRAPESR” for “CLN3_4_489aa” and “TLEGKKK” for “CLN3_235_181aa” are detected with protein Q-value < 0.05 in multiple samples (**Figure 6B**). The latter is particularly surprising given that “CLN3_235_181aa” belongs to the “major” disease-associated transcripts and they are predicted to undergo NMD. The identification of relevant peptides supports the translation of at least two transcripts in the human brain and whole blood. We further analysed the structures of these two protein isoforms and found that they show different orientations when compared with the canonical CLN3 protein based on AlphaFold 2 predictions: an inverted N-terminus in “CLN3_4_489aa” (**Figure 6D**) and an inverted C-terminus in “CLN3_235_181aa” (**Figure 6C**). These findings suggest potential differences in the functionality of these two protein isoforms as compared to canonical CLN3 protein.

**Figure 6.**
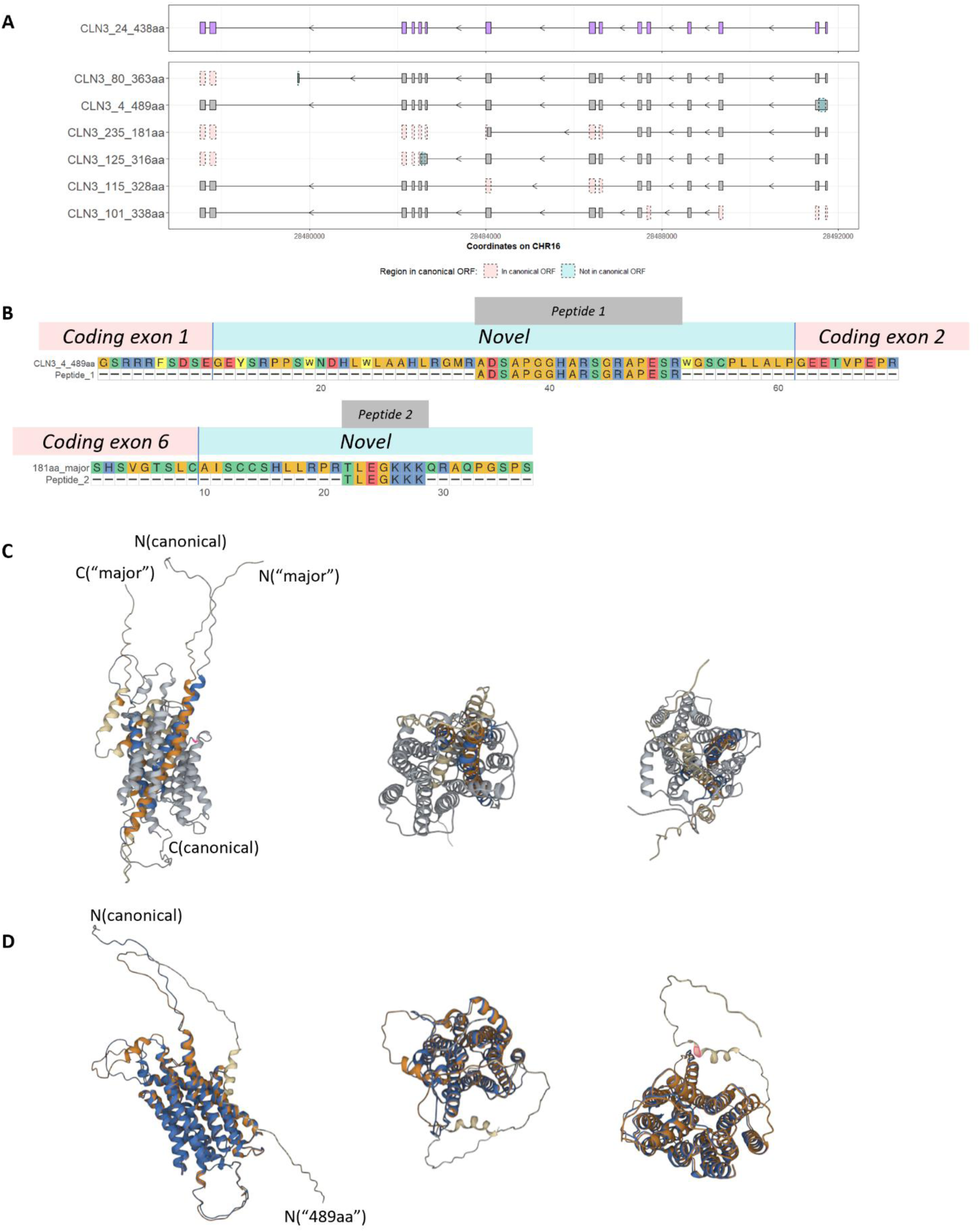
Validation of *CLN3* ORFs with unique sequences. Selected ORFs with unique sequences (CLN3_4_489aa, CLN3_125_316aa, CLN3_101_338aa, CLN3_81_363aa, CLN3_235_181aa, and CLN3_115_328aa) are compared with the canonical *CLN3* ORF “CLN3_24_438aa”. Grey boxes show exons of these ORFs, with pink boxes showing regions included in the canonical ORF but not in these ORFs and blue boxes showing regions included in these ORFs but not in the canonical ORF in **A**. In **B**, detected peptides “ADSAPGGHARSGRAPESR” for “CLN3_4_489aa” and “TLEGKKK” for “CLN3_235_181aa” are aligned to the corresponding ORFs, with amino acids that are not present in the canonical CLN3 protein shown as “Novel”. Protein products structures of “CLN3_24_438aa”, “CLN3_4_489aa” and “CLN3_235_181aa” (from the “major” transcripts) are predicted by AlphaFold 2.0 in **C** and **D**. The structure of the canonical CLN3 protein (blue and grey in **C** and **D**) is aligned against products of “CLN3_235_181aa” (orange and cream in **C**) and “CLN3_4_489aa” (orange and cream in **D**). Both protein isoforms show different orientations compared with the canonical 438aa CLN3 protein, “CLN3_235_181aa” has an inverted C-terminus and “CLN3_4_489aa” has an inverted N-terminus.

## Discussion

The *CLN3* transcription shows greater complexity than other genes. The number of transcripts for *CLN3* is 62 on Ensembl 110, which is much higher than the average of 3.42 transcripts per gene for GTEx V8 short-read RNA sequencing data (36). We detected 172 *CLN3* transcripts with a large number of unannotated *CLN3* transcripts (N = 147) in this study. Interestingly, only 30 out of the 64 annotated transcripts (GENCODE 29) are detected. There are 34 transcripts without support from ENCODE long-read RNA-seq data, in addition to the MANE select *CLN3* transcript ENST00000636147.2 in the current annotation (Ensembl 110, GENCODE 44). This MANE transcript does not match the most commonly used transcript (CLN3_24_438aa_5UTR_136_3UTR_49) or any other *CLN3* transcripts detected in our study. In fact, within our *CLN3* data, there is no dominantly expressed *CLN3* transcript. The most abundant *CLN3* transcript has median usage of 42.9% across 99 samples, well below the estimated usage of dominant transcripts of around 85% in human tissues and 75% in cell lines (34). In addition, the median usage of transcripts contain the canonical ORF is 66.7%, leaving around one-third of *CLN3* transcripts encoding non-canonical CLN3 protein isoforms. These findings highlight the importance of investigating the full variety of *CLN3* transcripts and studying *CLN3* in terms of highly used non-canonical transcripts and their products.

The large number of novel transcripts identified in this study can be partly explained by the fact that previous discoveries based on short-read RNA sequencing data would have been limited by the *CLN3-NPIPB7* readthrough gene, ENSG00000261832. ENSG00000261832 is annotated due to the existence of transcripts generated through the splicing together of *CLN3* and *NPIPB7* exons. This gene contains exons of both *CLN3* and *NPIPB7* and so will result in duplicated sequences in the reference, preventing the unique mapping of *CLN3* short-read sequencing data (33). These multi-mapped reads are typically removed in multiple sources including GTEx (16), IntroVerse (37), and recount3 (38). As a result, *CLN3* may have been both inaccurately quantified and annotated. Future long-read RNA sequencing studies for *CLN3* will need to consider this readthrough gene to ensure that the analysis pipeline does not duplicate or remove reads inappropriately.

The existence of *CLN3* transcripts with the same ORFs but different UTRs brings further complexity to *CLN3* annotation. The different transcription start sites detected in this study suggest the presence and use of alternative promoters for *CLN3*. Work is needed to investigate whether the usage of alternative promoters to generate transcripts with the same ORF but different 5’UTRs is cell-or tissue-specific (39), and whether regulatory elements, such as binding sites of RNA-binding proteins (RBPs), or secondary structures (40) are regulating translation efficiency (41) or mRNA stability (42). Most *CLN3* transcripts have closely located polyadenylation sites (PASs), however the 3’UTR of the most abundant *CLN3* transcript in our dataset has a distinct structure spanning 2164 nucleotides and is detected in all selected samples. Work is needed to investigate whether these different 3’UTRs serve to regulate gene expression levels by altering the mRNA localization, regulatory elements including micro-RNA and RBPs binding sites, or NMD status (43-48).

Relevant to understanding CLN3 disease is our observation of two specific *CLN3* transcripts previously thought to be generated only in disease states are detected in “healthy” control samples. Due to the rarity of juvenile CLN3 disease (49), it is extremely unlikely that all ENCODE tissue donors containing these disease-associated transcripts are carriers or patients. Instead, these transcripts are likely to be naturally occurring, but significantly increase in expression in patients with classic juvenile CLN3 disease. Previous research has indicated these translated disease-associated transcripts retain some functions and that they are not deleterious (7). The results of long-read RNA sequencing of patients with CLN3 disease are not yet available to allow comparison of the relative usage of transcripts from healthy tissues.

The existence of different CLN3 protein isoforms is supported by mass spectrometry data. Notably, all “major” disease-associated transcripts containing ORF “CLN3_253_181aa” are predicted to undergo NMD based on the distance between the most downstream exon-exon junction and the stop codons. It has been reported that transcripts with premature termination codons (PTCs) could escape NMD and produce truncated proteins (50, 51). The non-toxic truncated protein could partially rescue the disease phenotype (52). Thus, protein-level evidence will be needed to determine if a transcript with a PTC is NMD-sensitive.

Currently, cryo-electron microscopy or X-ray crystallography data for the canonical CLN3 protein is not available. Therefore, we are relying on AlphaFold topology predictions which show the canonical CLN3 protein has the N-terminus and C-terminus at different sides of the membrane (53). However, the topology predictions for “CLN3_253_181aa" and "CLN3_4_489aa" show the N-terminus and C-terminus of each protein isoform are on the same sides of the membrane. With the relative orientation of domains/motifs changing, the function of these isoforms could be different to the canonical CLN3 protein (54). “CLN3_253_181aa”, which has the first 153 amino acids of the canonical CLN3 protein and 28 non-canonical out-of-frame amino acids, is missing two lysosomal targeting motifs, ^242^EEE(X)8LI^254^, and ^409^M(X)9G^419^ of the canonical protein (55, 56). For “CLN3_4_489aa”, which has the N-terminus presenting at the opposite side as the N-terminus of the canonical CLN3 protein, the interaction with protein Rab7 which controls vesicular transport to late endosomes and lysosomes might be affected (57). Consequently, loss of the terminal portion of the protein (e.g., CLN3_253_181aa) or having an inverted terminus (e.g., CLN3_4_489aa) could lead to different trafficking and thus different functions compared with the canonical CLN3 protein. This identification of potentially structurally and functionally diverse protein isoforms provides us with future new perspectives from which to examine the biology of CLN3 and disease pathogenesis.

## Conclusion

Overall, this study expands our knowledge of *CLN3* transcription. We found that the overlapping *CLN3*-*NPIPB7* readthrough gene could affect accurate quantification and annotation of *CLN3.* With this information, we identified novel *CLN3* transcripts, as well as novel ORFs and UTRs, and characterised their usage. Our analysis reveals there is no dominantly expressed *CLN3* transcript and around one-third of transcripts encoding non-canonical CLN3 protein isoforms have not been studied before. We also identified transcripts that were thought to be expressed only in juvenile CLN3 disease patients in healthy controls. Additionally, alternative UTRs suggest potential regulatory roles which could affect CLN3 protein abundance. Utilising mass spectrometry data to validate the detection of transcripts/ORFs is a powerful tool. The detection of peptides belonging to non-canonical CLN3 protein isoforms demonstrates the translational potential of corresponding transcripts/ORFs and may have functional biological relevance. This highlights the importance of investigating non-canonical protein isoforms in detail to further our understanding of their functions and their role in disease pathogenesis. The insights gained from this study have implications for studying patient samples in the future, as they provide a valuable reference when examining how the "1-kb" deletion affects the *CLN3* transcription and contributes to disease pathogenesis.

## Abbreviations

NCLs: Neuronal ceroid lipofuscinoses
MFS: major facilitator superfamily
ORFs: open-reading frames
UTRs: untranslated regions
TPM: transcripts per million
RPK: reads per kilobase
ID: identifier
CDS: coding sequence
NMD: nonsense-mediated decay
TSS: transcription start site
PAS: polyadenylation site
polyA: polyadenylation
MSA: multiple system atrophy
NSCLC: non-small-cell lung cancer
RBP: RNA-binding protein
PTC: premature termination codon

## Declarations

### Ethics approval and consent to participate

Not applicable

### Consent for publication

Not applicable

### Availability of data and materials

Comprehensive gene annotation of GENCODE release 29 (GENCODE 29) was downloaded from https://www.gencodegenes.org/human/release_29.html. Gene TPMs GTEx_Analysis_2017-06-05_v8_RNASeQCv1.1.9_gene_reads.gct.gz and exon-exon junction read counts GTEx_Analysis_2017- 06-05_v8_STARv2.5.3a_junctions.gct.gz of GTEx V8 can be accessed from GTEx portal (https://gtexportal.org/home/datasets/). The ENCODE long-read RNA sequencing data used in this paper was downloaded from ENCODE portal (https://www.encodeproject.org/). Public mass spectrometry datasets PXD026370 and PXD028605 can be downloaded from ProteomeXchange (https://www.proteomexchange.org/). Codes used to analyse the data and produce figures are accessible on GitHub (https://github.com/HYzhang800/CLN3_public_long_read).

### Authors’ Contributions

Conceptualisation: C.M., M.R. and S.E.M; Methodology: H.Y.Z., C.M., E.G., and M.R.; Data acquisition and analysis: H.Y.Z.; Data interpretation: H.Y.Z., C.M., E.G., M.R., and S.E.M.; Writing – original draft: H.Y.Z.; Writing – review & editing: C.M., E.G., M.R. and S.E.M.; Visualization: H.Y.Z.; Supervision: M.R. and S.E.M.; Project administration: C.M. and S.E.M.; Funding acquisition: C.M. M.R and S.E.M. All authors read and approved the final manuscript.

## Acknowledgements

Not applicable

