## supplemental_tables for "*CLN3* transcript complexity revealed by long-read RNA sequencing analysis"

**Table S1. Summary of five splice junctions showing the “readthrough” event in which exons from *CLN3* and *NPIPB7* are joined.**

This table shows five different splice junctions with splice donors in *CLN3* and splice acceptors in *NPIPB7.* For each splice junction and tissue, average read counts across all donors and tissue subtypes are summarised. The detection rates for each splice junction show the proportion of tissue donors in which specific junctions could be detected.

| Junction No. | Chr | Acceptor | Donor | Minimum  reads | Mean  reads | Maximum  reads | Tissues with the  maximum reads | Maximum detection rates | Tissues with the maximum detection rates  (N ≥ 100) |
| --- | --- | --- | --- | --- | --- | --- | --- | --- | --- |
| 1 | chr16 | 28466903 | 28476250 | 1.00 | 1.30 | 2.00 | Fallopian Tube | 76.1% | Brain |
| 2 | chr16 | 28466903 | 28482104 | 1.00 | 1.30 | 1.87 | Pituitary | 63.2% | Brain |
| 3 | chr16 | 28466903 | 28477463 | 1.34 | 2.12 | 3.66 | Thyroid | 97.8% | Brain |
| 4 | chr16 | 28471175 | 28476250 | 1.00 | 1.23 | 1.68 | Brain | 83.9% | Brain |
| 5 | chr16 | 28466903 | 28477011 | 1.00 | 1.12 | 1.50 | Kidney | 24.7% | Brain |

**Table S2. Summary of selected ENCODE samples.**

| Experiment. accession | Tissue | Type | Description | ENCODE. Organ |
| --- | --- | --- | --- | --- |
| ENCSR398SKD | HL-60 | cell line | NA | Blood |
| ENCSR546DFO | HL-60 | cell line | NA | Blood |
| ENCSR121FDE | HL-60 | cell line | NA | Blood |
| ENCSR159ICU | HL-60 | cell line | NA | Blood |
| ENCSR930GRQ | HL-60 | cell line | NA | Blood |
| ENCSR583KAF | HL-60 | cell line | NA | Blood |
| ENCSR278ZPI | HL-60 | cell line | NA | Blood |
| ENCSR887LTD | HL-60 | cell line | NA | Blood |
| ENCSR700EBI | aorta | tissue | Homo sapiens aorta tissue female adult (41 years) | Blood vessel |
| ENCSR853YZN | posterior vena cava | tissue | Homo sapiens posterior vena cava tissue female adult (47 years) | Blood vessel |
| ENCSR687RWI | endothelial cell of umbilical vein | primary cell | NA | Blood vessel |
| ENCSR425HFS | aorta | tissue | Homo sapiens aorta tissue female adult (59 years) | Blood vessel |
| ENCSR138TAS | posterior vena cava | tissue | Homo sapiens posterior vena cava tissue female adult (59 years) | Blood vessel |
| ENCSR697ASE | middle frontal area 46 | tissue | Homo sapiens with Alzheimer's disease; dorsolateral prefrontal cortex tissue female adult (90 or above years) | Brain |
| ENCSR205QMF | middle frontal area 46 | tissue | Homo sapiens dorsolateral prefrontal cortex tissue female adult (88 years) | Brain |
| ENCSR169YNI | middle frontal area 46 | tissue | Homo sapiens dorsolateral prefrontal cortex tissue female adult (90 or above years) | Brain |
| ENCSR094NFM | middle frontal area 46 | tissue | Homo sapiens dorsolateral prefrontal cortex tissue female adult (90 or above years) | Brain |
| ENCSR462COR | middle frontal area 46 | tissue | Homo sapiens with Alzheimer's disease; dorsolateral prefrontal cortex tissue female adult (90 or above years) | Brain |
| ENCSR463IDK | middle frontal area 46 | tissue | Homo sapiens dorsolateral prefrontal cortex tissue female adult (79 years) | Brain |
| ENCSR316ZTD | middle frontal area 46 | tissue | Homo sapiens with Alzheimer's disease; dorsolateral prefrontal cortex tissue female adult (90 or above years) | Brain |
| ENCSR257YUB | middle frontal area 46 | tissue | Homo sapiens with Alzheimer's disease; dorsolateral prefrontal cortex tissue male adult (90 or above years) | Brain |
| ENCSR690QHM | middle frontal area 46 | tissue | Homo sapiens with Alzheimer's disease; dorsolateral prefrontal cortex tissue female adult (90 or above years) | Brain |
| ENCSR309IKK | WTC11 | cell line | NA | Connective tissue, Skin |
| ENCSR507JOF | WTC11 | cell line | NA | Connective tissue, Skin |
| ENCSR470HYQ | chondrocyte | In vitro differentiated cells | NA | Connective tissue |
| ENCSR296PQZ | osteocyte | in vitro differentiated cells | NA | Connective tissue |
| ENCSR113MEY | IMR-90 | cell line | NA | Connective tissue |
| ENCSR648QZP | mesenteric fat pad | tissue | Homo sapiens mesenteric fat pad tissue female adult (59 years) | Connective tissue |
| ENCSR902GAF | HFFc6 | cell line | NA | Connective tissue |
| ENCSR127HKN | endodermal cell | in vitro differentiated cells | NA | Embryo |
| ENCSR319VGI | H1 | cell line | NA | Embryo |
| ENCSR271KEJ | H1 | cell line | NA | Embryo |
| ENCSR972ABM | endodermal cell | in vitro differentiated cells | NA | Embryo |
| ENCSR044ARQ | H9 | cell line | NA | Embryo |
| ENCSR056MYH | neural crest cell | in vitro differentiated cells | NA | Embryo |
| ENCSR904KAT | endodermal cell | in vitro differentiated cells | NA | Embryo |
| ENCSR563RLX | adrenal gland | tissue | Homo sapiens adrenal gland tissue female adult (41 years) | Endocrine gland |
| ENCSR995WKW | adrenal gland | tissue | Homo sapiens adrenal gland tissue male adult (54 years) | Endocrine gland |
| ENCSR530BOC | progenitor cell of endocrine pancreas | in vitro differentiated cells | NA | Endocrine gland |
| ENCSR130CIK | type B pancreatic cell | in vitro differentiated cells | NA | Endocrine gland |
| ENCSR657HJW | right lobe of liver | tissue | Homo sapiens right lobe of liver tissue female adult (41 years) | Endocrine gland |
| ENCSR081NRO | adrenal gland | tissue | Homo sapiens adrenal gland tissue male adult (37 years) | Endocrine gland |
| ENCSR293MOX | right lobe of liver | tissue | Homo sapiens right lobe of liver tissue female child (16 years) | Endocrine gland |
| ENCSR899GAP | Right ventricle myocardium superior | tissue | Homo sapiens Right ventricle myocardium superior tissue male adult (60 years) | Heart |
| ENCSR549ELD | cardiac septum | tissue | Homo sapiens cardiac septum tissue female adult (41 years) | Heart |
| ENCSR435UUS | right cardiac atrium | tissue | Homo sapiens right cardiac atrium tissue male adult (60 years) | Heart |
| ENCSR777CCI | left ventricle myocardium superior | tissue | Homo sapiens left ventricle myocardium superior tissue male adult (60 years) | Heart |
| ENCSR782LGT | heart right ventricle | tissue | Homo sapiens heart right ventricle tissue female adult (46 years) | Heart |
| ENCSR984OAE | heart right ventricle | tissue | Homo sapiens heart right ventricle tissue male adult (40 years) | Heart |
| ENCSR514YQN | right cardiac atrium | tissue | Homo sapiens right cardiac atrium tissue female adult (46 years) | Heart |
| ENCSR728TXV | right cardiac atrium | tissue | Homo sapiens right cardiac atrium tissue male adult (40 years) | Heart |
| ENCSR994YZY | heart left ventricle | tissue | Homo sapiens heart left ventricle tissue male adult (40 years) | Heart |
| ENCSR786FLO | left ventricle myocardium inferior | tissue | Homo sapiens left ventricle myocardium inferior tissue male adult (60 years) | Heart |
| ENCSR591OZR | Right ventricle myocardium inferior | tissue | Homo sapiens Right ventricle myocardium inferior tissue male adult (60 years) | Heart |
| ENCSR700XDQ | heart left ventricle | tissue | Homo sapiens heart left ventricle tissue female adult (46 years) | Heart |
| ENCSR553SVP | right cardiac atrium | tissue | Homo sapiens right cardiac atrium tissue female adult (59 years) | Heart |
| ENCSR329ZQG | heart right ventricle | tissue | Homo sapiens heart right ventricle tissue female adult (59 years) | Heart |
| ENCSR194YUY | heart left ventricle | tissue | Homo sapiens heart left ventricle tissue female adult (53 years) | Heart |
| ENCSR575LWI | heart left ventricle | tissue | Homo sapiens heart left ventricle tissue female adult (59 years) | Heart |
| ENCSR424QFN | left cardiac atrium | tissue | Homo sapiens left cardiac atrium tissue female adult (59 years) | Heart |
| ENCSR323XND | lower lobe of right lung | tissue | Homo sapiens lower lobe of right lung tissue male adult (60 years) | Lung |
| ENCSR096QUP | upper lobe of right lung | tissue | Homo sapiens upper lobe of right lung tissue male adult (60 years) | Lung |
| ENCSR986WKB | lower lobe of left lung | tissue | Homo sapiens lower lobe of left lung tissue male adult (60 years) | Lung |
| ENCSR426KOP | left lung | tissue | Homo sapiens left lung tissue male adult (40 years) | Lung |
| ENCSR746ITG | left lung | tissue | Homo sapiens left lung tissue female child (16 years) | Lung |
| ENCSR261GOA | lower lobe of left lung | tissue | Homo sapiens lower lobe of left lung tissue female adult (59 years) | Lung |
| ENCSR676IWT | GM23338 | cell line | NA | Skin |

**Table S3. Expression level of *CLN3*-*NPIPB7* readthrough gene (ENSG00000261832) across all ENCODE-defined organs.**

| Organs | Samples in total | Samples detected | Detection rates | Min TPM | Median TPM | Max TPM |
| --- | --- | --- | --- | --- | --- | --- |
| Blood | 16 | 13 | 81.3% | 0.13 | 0.49 | 1.56 |
| Blood vessel | 6 | 3 | 50.0% | 0.09 | 0.57 | 0.72 |
| Brain | 9 | 7 | 77.8% | 0.16 | 0.39 | 1.43 |
| Connective tissue | 16 | 11 | 68.8% | 0.10 | 0.30 | 1.39 |
| Embryo | 18 | 10 | 55.6% | 0.15 | 0.38 | 0.55 |
| Endocrine gland | 9 | 7 | 77.8% | 0.19 | 0.46 | 0.84 |
| Heart | 17 | 6 | 35.3% | 0.16 | 0.26 | 0.42 |
| Lung | 6 | 6 | 100.0% | 0.13 | 0.50 | 1.60 |
| Skin | 8 | 3 | 37.5% | 0.10 | 0.15 | 0.17 |

**Table S4. Summary of tissue-specific transcripts.**

| ID | Tissue | ENCODE Organ | Usage |
| --- | --- | --- | --- |
| CLN3_4_489aa_5UTR_60_3UTR_52 | posterior vena cava | blood vessel | 25.0% |
| CLN3_267_109aa_5UTR_169_3UTR_14 | right lobe of liver | endocrine gland | 13.3% |
| CLN3_235_181aa_5UTR_132_3UTR_98 | heart left ventricle | heart | 10.0% |
| CLN3_235_181aa_5UTR_no_3UTR_108 | heart left ventricle | heart | 10.0% |
| CLN3_119_321aa_5UTR_132_3UTR_72 | right lobe of liver | endocrine gland | 9.1% |
| CLN3_24_438aa_5UTR_194_3UTR_52 | posterior vena cava | blood vessel | 8.3% |
| CLN3_101_338aa_5UTR_128_3UTR_52 | middle frontal area 46 | brain | 8.1% |
| CLN3_91_355aa_5UTR_132_3UTR_52 | GM23338 | skin | 8.0% |
| CLN3_207_223aa_5UTR_29_3UTR_52 | GM23338 | skin | 6.7% |
| CLN3_166_267aa_5UTR_no_3UTR_71 | heart right ventricle | heart | 6.3% |
| CLN3_171_262aa_5UTR_no_3UTR_86 | heart right ventricle | heart | 6.3% |
| CLN3_24_438aa_5UTR_195_3UTR_52 | right cardiac atrium | heart | 6.3% |
| CLN3_57_384aa_5UTR_no_3UTR_44 | left ventricle myocardium inferior | heart | 5.6% |
| CLN3_205_226aa_5UTR_132_3UTR_16 | middle frontal area 46 | brain | 5.0% |
| CLN3_166_267aa_5UTR_17_3UTR_71 | H1 | embryo | 4.5% |
| CLN3_22_440aa_5UTR_65_3UTR_52 | right cardiac atrium | heart | 4.5% |
| CLN3_24_438aa_5UTR_215_3UTR_52 | right lobe of liver | endocrine gland | 4.5% |
| CLN3_41_421aa_5UTR_no_3UTR_54 | H1 | embryo | 4.5% |
| CLN3_77_366aa_5UTR_128_3UTR_52 | right lobe of liver | endocrine gland | 4.5% |
| CLN3_119_321aa_5UTR_65_3UTR_71 | heart right ventricle | heart | 4.3% |
| CLN3_24_438aa_5UTR_61_3UTR_44 | heart right ventricle | heart | 4.3% |
| CLN3_252_146aa_5UTR_122_3UTR_52 | middle frontal area 46 | brain | 4.0% |
| CLN3_273_55aa_5UTR_137_3UTR_36 | GM23338 | skin | 4.0% |
| CLN3_76_366aa_5UTR_52_3UTR_52 | WTC11 | connective tissue | 3.8% |
| CLN3_24_438aa_5UTR_134_3UTR_44 | middle frontal area 46 | brain | 3.6% |
| CLN3_26_437aa_5UTR_65_3UTR_52 | left cardiac atrium | heart | 3.3% |
| CLN3_101_338aa_5UTR_296_3UTR_52 | WTC11 | connective tissue | 3.2% |
| CLN3_23_440aa_5UTR_132_3UTR_52 | WTC11 | connective tissue | 3.2% |
| CLN3_24_438aa_5UTR_136_3UTR_44 | aorta | blood vessel | 3.2% |
| CLN3_119_321aa_5UTR_132_3UTR_70 | endodermal cell | embryo | 3.0% |
| CLN3_235_181aa_5UTR_134_3UTR_107 | Right ventricle myocardium superior | heart | 2.9% |
| CLN3_135_307aa_5UTR_132_3UTR_75 | heart right ventricle | heart | 2.9% |
| CLN3_22_440aa_5UTR_64_3UTR_52 | heart right ventricle | heart | 2.9% |
| CLN3_252_146aa_5UTR_262_3UTR_52 | heart right ventricle | heart | 2.9% |
| CLN3_119_321aa_5UTR_132_3UTR_45 | endodermal cell | embryo | 2.4% |
| CLN3_132_309aa_5UTR_92_3UTR_63 | right cardiac atrium | heart | 2.4% |
| CLN3_153_285aa_5UTR_18_3UTR_52 | H1 | embryo | 2.4% |
| CLN3_96_344aa_5UTR_64_3UTR_127 | H1 | embryo | 2.4% |
| CLN3_113_331aa_5UTR_303_3UTR_79 | middle frontal area 46 | brain | 2.4% |
| CLN3_235_181aa_5UTR_132_3UTR_110 | middle frontal area 46 | brain | 2.4% |
| CLN3_209_221aa_5UTR_52_3UTR_71 | progenitor cell of endocrine pancreas | endocrine gland | 2.3% |
| CLN3_177_254aa_5UTR_64_3UTR_52 | H1 | embryo | 2.3% |
| CLN3_261_127aa_5UTR_92_3UTR_110 | endodermal cell | embryo | 2.2% |
| CLN3_207_223aa_5UTR_104_3UTR_52 | HFFc6 | connective tissue | 1.9% |
| CLN3_207_223aa_5UTR_101_3UTR_52 | progenitor cell of endocrine pancreas | endocrine gland | 1.8% |
| CLN3_92_355aa_5UTR_132_3UTR_63 | middle frontal area 46 | brain | 1.7% |
| CLN3_80_363aa_5UTR_64_3UTR_63 | WTC11 | connective tissue | 1.7% |
| CLN3_266_115aa_5UTR_237_3UTR_52 | HL-60 | HL60 | 1.6% |
| CLN3_14_453aa_5UTR_78_3UTR_52 | middle frontal area 46 | brain | 1.5% |
| CLN3_275_48aa_5UTR_319_3UTR_76 | middle frontal area 46 | brain | 1.5% |
| CLN3_20_441aa_5UTR_68_3UTR_52 | endodermal cell | embryo | 1.5% |
| CLN3_24_438aa_5UTR_196_3UTR_52 | lower lobe of right lung | lung | 1.5% |
| CLN3_216_216aa_5UTR_128_3UTR_79 | endodermal cell | embryo | 1.1% |
| CLN3_107_336aa_5UTR_16_3UTR_55 | WTC11 | connective tissue | 1.0% |
| CLN3_207_223aa_5UTR_273_3UTR_52 | HL-60 | HL60 | 1.0% |
| CLN3_171_262aa_5UTR_no_3UTR_88 | HL-60 | HL60 | 1.0% |
| CLN3_249_148aa_5UTR_246_3UTR_63 | HL-60 | HL60 | 0.9% |
| CLN3_119_321aa_5UTR_64_3UTR_70 | progenitor cell of endocrine pancreas | endocrine gland | 0.9% |
| CLN3_119_321aa_5UTR_158_3UTR_71 | HL-60 | HL60 | 0.8% |
| CLN3_171_262aa_5UTR_no_3UTR_79 | HL-60 | HL60 | 0.7% |
| CLN3_132_309aa_5UTR_283_3UTR_63 | HL-60 | HL60 | 0.7% |
| CLN3_166_267aa_5UTR_283_3UTR_70 | HL-60 | HL60 | 0.6% |
| CLN3_252_146aa_5UTR_259_3UTR_52 | HL-60 | HL60 | 0.6% |
| CLN3_125_316aa_5UTR_146_3UTR_79 | HL-60 | HL60 | 0.6% |
| CLN3_252_146aa_5UTR_210_3UTR_52 | HL-60 | HL60 | 0.6% |
| CLN3_252_146aa_5UTR_244_3UTR_52 | HL-60 | HL60 | 0.6% |
| CLN3_80_363aa_5UTR_146_3UTR_63 | HL-60 | HL60 | 0.5% |
| CLN3_24_438aa_5UTR_65_3UTR_44 | HL-60 | HL60 | 0.5% |
| CLN3_80_363aa_5UTR_158_3UTR_63 | HL-60 | HL60 | 0.5% |
| CLN3_24_438aa_5UTR_206_3UTR_52 | HL-60 | HL60 | 0.5% |
| CLN3_24_438aa_5UTR_143_3UTR_52 | HL-60 | HL60 | 0.5% |
| CLN3_249_148aa_5UTR_265_3UTR_63 | HL-60 | HL60 | 0.4% |
| CLN3_24_438aa_5UTR_190_3UTR_52 | HL-60 | HL60 | 0.3% |
| CLN3_24_438aa_5UTR_164_3UTR_52 | HL-60 | HL60 | 0.3% |
| CLN3_24_438aa_5UTR_184_3UTR_52 | HL-60 | HL60 | 0.3% |
