## supplemental_figures for "*CLN3* transcript complexity revealed by long-read RNA sequencing analysis"

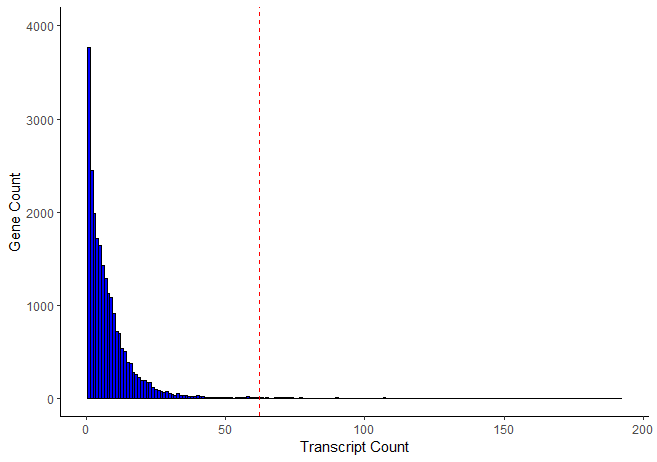

**Figure S1. Distribution of the number of transcripts per gene in Ensembl 110.**

This plot shows the distribution of the number of transcripts per gene in Ensembl 110. The red dashed line marks the position of CLN3.

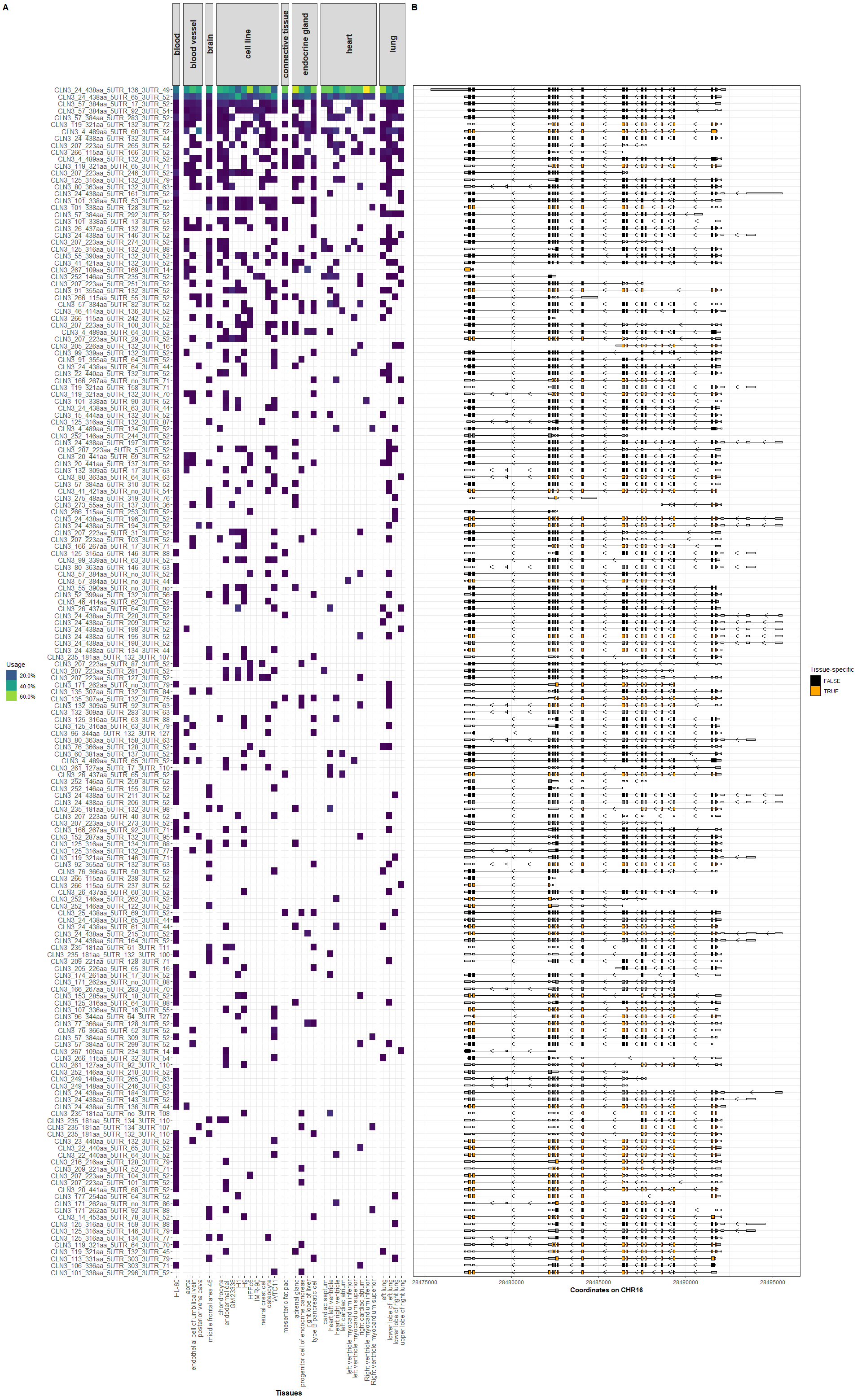

**Figure S2. Usage and tissue-specificity for all 172 valid *CLN3* transcripts.**

Usage of valid transcripts across tissue types is summarised and plotted using heat maps with yellow showing high usage and dark blue showing low usage. Structures of all 172 valid transcripts are plotted, with ORFs coloured by tissue-specificity. Orange shows tissue-specific transcripts and black shows non-tissue-specific transcripts. UTRs are represented by shorter grey boxes.
